# The chloride cotransporter NKCC1 regulates self-renewal of hippocampal neural stem cells via the transcription factor Sox11

**DOI:** 10.1101/2025.07.01.662334

**Authors:** Antonia Blank, Sneha Adhikarla, Julius Broesske, Lucas Kaiser, Magdalene Rippe, Madlen Haase, Marco Groth, Gregor Stein, Tina Andrä, Jessica Mesterheide, Anna-Lena Fleischer, Tom Flossmann, Iris Schäffner, Elisabeth Sock, Knut Kirmse, Christian A. Hübner, Knut Holthoff, Christian Schmeer, Dieter Chichung Lie, Silke Keiner

## Abstract

The GABAergic-mediated depolarization plays a key role in controlling stem cell fate and neurogenesis within the dentate gyrus of the hippocampal formation. This depolarization effect is highly dependent on the balance between the chloride co-transporters NKCC1 and KCC2. It is not known how changes in NKCC1 modulate the fate of Nestin-positive stem cells (NSCs) in the hippocampus during adult neurogenesis. In our study, we demonstrate that a knockout of *Nkcc1* in NSCs increase their proliferation by symmetric self-renewal and expand the stem cell pool. Using single-cell RNA sequencing, we identified Sox11 as a key transcription factor that is significantly downregulated following *Nkcc1* knockout. In agreement with this finding, we found that *Sox11* knockout enhances proliferation and self-renewal of NSCs, which is also marked by an increase in symmetric stem cell division. Based on these findings, we propose that altering *Nkcc1* expression in NSCs shifts their fate from neurogenesis towards self-renewal via Sox11 regulation. Furthermore, we observed that NKCC1 levels decline in NSCs during aging, which correlates with a further increase in self-renewal. Our data strongly suggest that the age-related decline in NKCC1 levels promote symmetric division and self-renewal contributing to the age-dependent decrease in neurogenesis. NKCC1 via Sox11 is a key regulator of NSCs fate decision, critically balancing self-renewal and neuronal differentiation in the adult hippocampus.

**Highlights:** NKCC1 is a key factor in regulating the Nestin-positive neural stem cell symmetric self-renewal.

Sox11 is a downstream effector of NKCC1 in Nestin-positive stem cells and is involved in expansion of the stem cell pool.

Aging reduces NKCC1 expression in Nestin-positive stem cells and increases their self-renewal.

**Graphical abstract:** 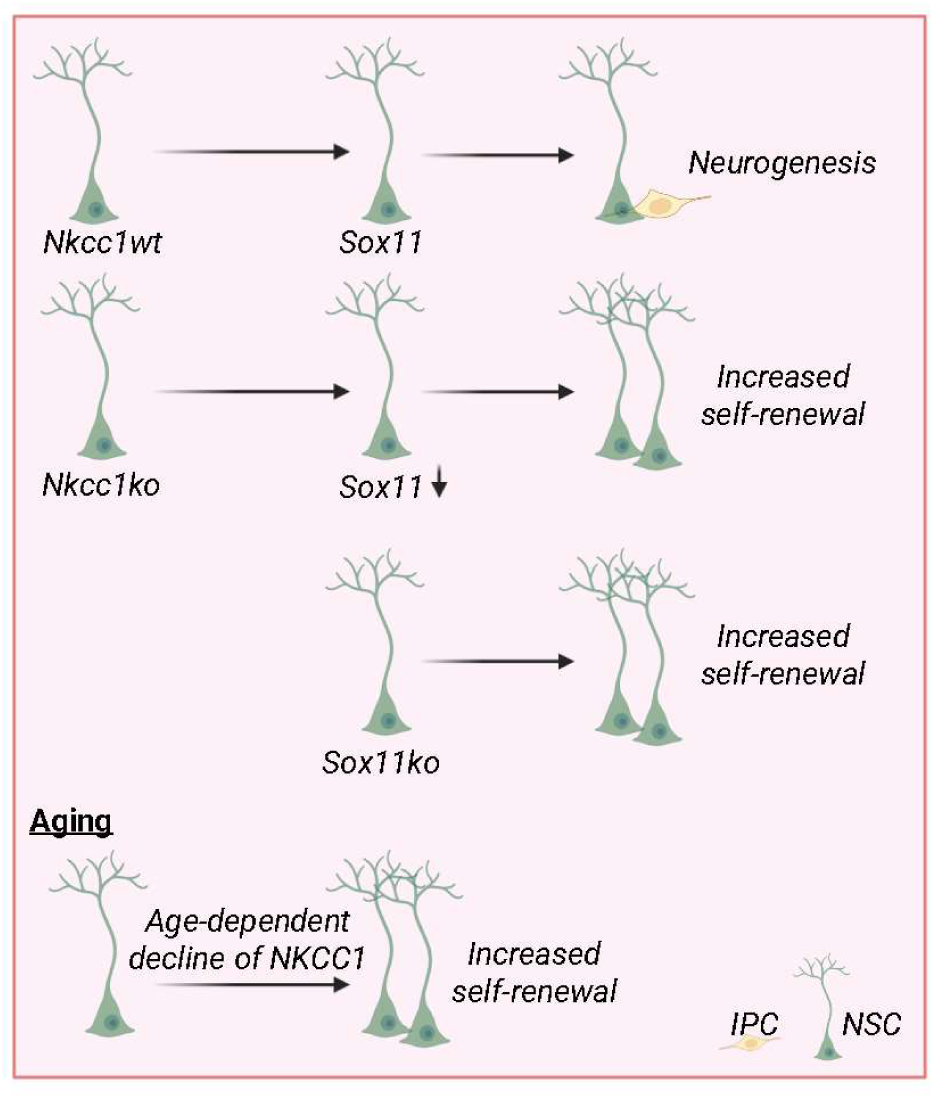

## Introduction

Activation and self-renewal of NSCs are essential processes for maintaining neurogenesis in the adult brain. In the hippocampus, NSCs exist in a quiescent state, where they do not actively proliferate but can be stimulated to divide triggered by various extrinsic and intrinsic signals (Abbott & Nigussie, 2020; Urban *et al*, 2019). A controlled activation of these NSCs is crucial to maintain a balance between self-renewal and differentiation, ensuring long-term regeneration of neuronal networks (Bonaguidi *et al*, 2011). Activation of NSCs is induced by multiple signals, including neuronal activity, growth factors, and neurotransmitters. Notably, GABA (γ-aminobutyric acid) plays a critical role in NSCs activation, by inducing depolarization of NSCs via GABA_A_ receptors (Bao *et al*, 2017; Song *et al*, 2012; Zhang *et al*, 2023). It has been shown that alterations in GABA_A_ receptor function promote proliferation and symmetric NSC division (Song *et al*., 2012). The effect of GABA through the GABA_A_ receptor depends on intracellular chloride homeostasis, which is regulated by the chloride cotransporters NKCC1 and KCC2. NKCC1 (Na-K-2Cl-Cotransporter 1, encoded by the gene slc12a2) maintains a high intracellular chloride concentration thus enhancing GABA-induced depolarization, whereas KCC2 (K-Cl-Cotransporter 2), predominantly expressed in mature neurons, promotes hyperpolarization by lowering intracellular chloride concentration (Watanabe & Fukuda, 2015; Zhang *et al*, 2021). Self-renewal of NSCs is crucial for sustaining the NSC pool over time. NSCs undergo a finely tuned balance between asymmetric and symmetric cell division in order to maintain adult neurogenesis (Ibrayeva *et al*, 2021). In particular, NSCs are essential for NSC maintenance, as they have the ability to self-renew and give rise to neurons and astrocytes throughout life (Bonaguidi *et al*., 2011; Ibrayeva *et al*., 2021; Pilz *et al*, 2018). Adult mammalian NCSs and neurogenesis are tightly regulated by cell extrinsic and intrinsic factors (Matsubara *et al*, 2021). One of the main intrinsic mechanisms involves the control of cellular processes by transcriptional regulation of gene expression. A key regulator of neurogenesis is the transcription factor Sox11, both during embryonic development and in adulthood (Haslinger *et al*, 2009; Mu *et al*, 2012; Wang *et al*, 2013). During embryogenesis, the ablation of *Sox11* disrupts neural progenitor cell (NPC) proliferation, neuronal migration, and differentiation. In adult brains, the loss of *Sox11* in NPCs impairs hippocampal neurogenesis. Using a retroviral approach, it was demonstrated that the ablation of both *Sox11* and the closely related *Sox4* inhibits neurogenesis from adult NPCs *in vitro* and *in vivo* (Mu *et al*., 2012).

With aging, hippocampal neurogenesis declines, accompanied by changes in NSC proliferation and an increase in symmetric NSC division (Ibrayeva *et al*., 2021; Wu *et al*, 2023). The mechanisms leading to fate changes and the role of the chloride cotransporter NKCC1 in this process remain unclear. In the present study, we knocked out *Nkcc1* in NSCs and observed increased NSC proliferation, mediated by an increase in symmetric cell division. As a downstream mechanism, we identified decreased expression of *Sox11*, following *Nkcc1* knockout. The *Sox11* knockout in NSCs also led to increased proliferation and enlarged the NSC pool. NKCC1 levels during aging were decreased and was associated with an increased NSCs self-renewal, similar as observed after the *Nkcc1* knockout. This suggests a mechanism involving reduced NKCC1 levels and increased symmetric NSC division in aging, leading to impaired generation of new neurons.

## RESULTS

### *Nkcc1* knockout promotes stem cell proliferation and symmetric division

To investigate the influence of NKCC1 on the proliferative properties of NSCs, we used 2-month-old genetically modified *Nestin-CreER^T2^/tdTomato* as controls and *Nestin-CreER^T2^/Nkcc1^fl/fl^/tdTomato* mice expressing the fluorescent reporter protein tdTomato in NSCs. After tamoxifen delivery to deplete *Nkcc1*, we analyzed various endogenous proliferation markers such as MCM2, as well as labeling of proliferative cells via the Thymidine-analogues CldU and IdU, which were injected intraperitoneally. We identified NSCs in the subgranular zone of the dentate gyrus by their typical morphological appearance (apical processes reaching into the molecular layer and the triangular soma), which was visualized through their tdTomato expression and characteristic markers such as GFAP and *Sox*2. NPCs were identified by means of the markers TBR2 and DCX.

To characterize the NKCC1-dependent changes in NSC proliferation we used the thymidine analogues CldU and IdU after tamoxifen injection. These markers are incorporated into the DNA during the S-phase of the cell cycle and are subsequently passed on to the resulting progeny (Fig. 1A) (Encinas *et al*, 2011). In both the control and *Nkcc1* knockout (*Nkcc1ko)* groups, IdU injection was administered either 16 or 24 hours (h) after the CIdU injection (Fig. 1A). We find that the proportion of CldU-positive NSCs among all tdTomato-positive cells significantly increases in *Nkcc1ko* mice compared to controls 24h after injection (Fig. 1B/C). Furthermore, the proliferation rate of IdU-positive NSCs is significantly increased at 16 to 24h. To analyze how many NSCs re-enter the cell cycle, the number of CldU-IdU-positive NSCs was determined 16h and 24h after CldU injection. A significant increase in NSC re-entry was observed in *Nkcc1ko* mice. Overall, the data suggest that *Nkcc1ko* in NSCs promotes proliferation and facilitates re-entry of previously proliferating NSCs into the cell cycle. To verify our findings, we administered a single low-dose of tamoxifen, which labeled a small number of tdTomato-positive NSCs in the dentate gyrus as previously described by (Bonaguidi *et al*., 2011). Animals were then scarified at different time points. Since the NSCs divide in place, it was possible to analyze clusters formed from NSCs over time in the different groups. Analyses of MCM2 expression showed increased proliferation within the clusters containing only NSCs in *Nkcc1ko* mice compared to controls (Fig. 1D-F). To better track the fate of the NSCs, we quantified the cluster compositions to characterize the NSC clonal progeny at 14- and 28-days post injection (Fig. 1G). This revealed that, particularly after *Nkcc1* knockout, the proportion of NSCs significantly increased compared to the control, at the expense of neuronal precursor cells (Fig. 1H/I).

**Figure 1.**
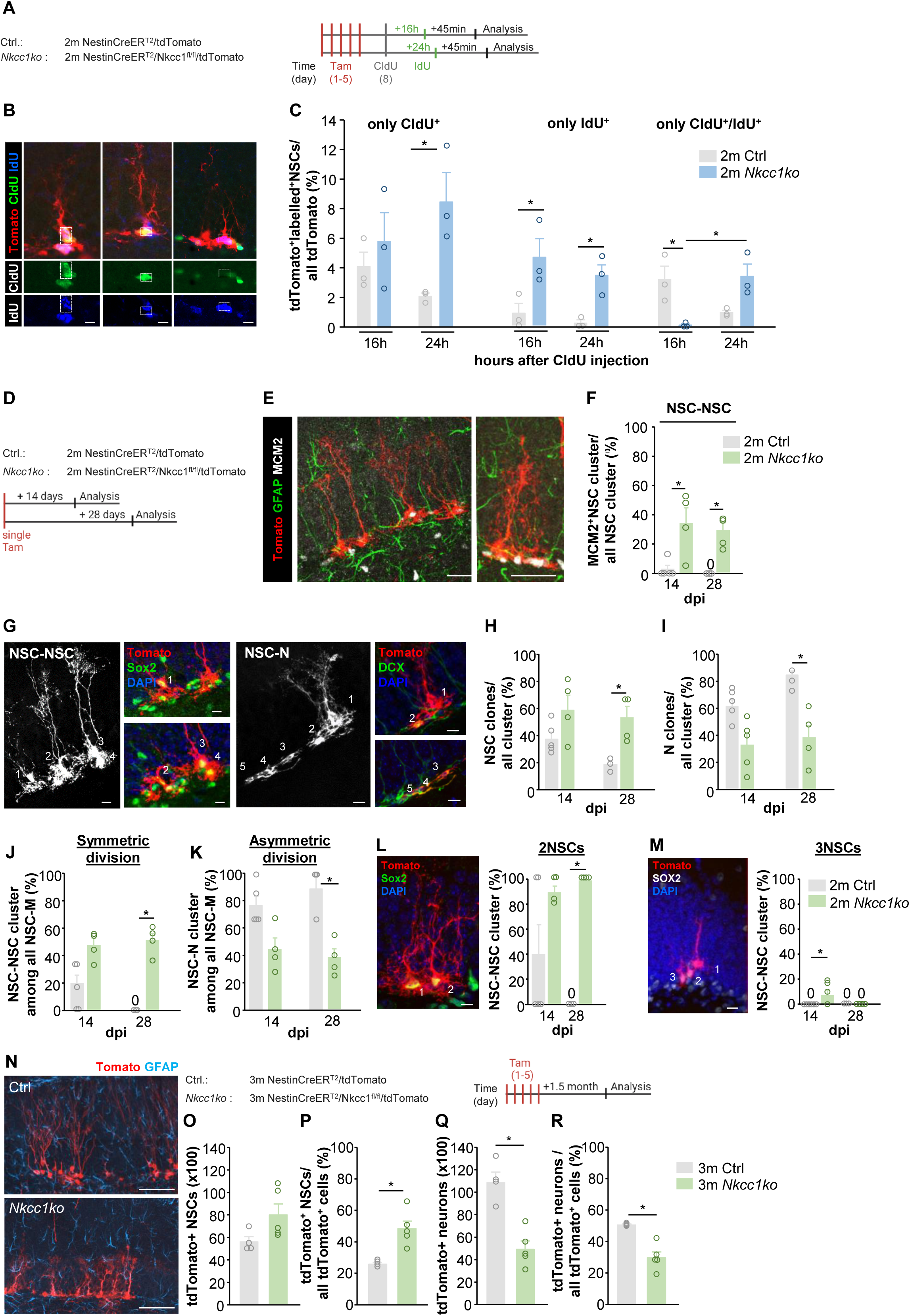
*Nkcc1* knockout induces NSC proliferation and self-renewal (A) To estimate proliferation of NSCs after *Nkcc1ko,* mice were injected with tamoxifen for 5 days followed by CldU (green) and IdU (blue) injections according to the scheme. IdU/CldU double-labeled NSCs are the ones that re-enter the cell cycle. 2-month-old *NestinCreER^T2^/tdTomato* (n=3) and *NestinCreER^T2^/Nkcc1^fl/fl^/tdTomato* (n=3) mice were used. (B) Representative confocal immunofluorescence images from the dentate gyrus showing NSCs co-labelled with the proliferation markers CldU and IdU. Scale bar: 10μm. (C) Quantification of NSCs positive for IdU, CldU or both (CIdU/IdU). Values represent mean ± SEM, Kruskal-Wallis-Test with posthoc Dunn’s Test, ∗p<0.05. (D) In vivo clonal tracing analysis of Nestin^+^ NSCs. Single tamoxifen injection was applied in 2-month-old *NestinCreER^T2^/tdTomato* mice and *NestinCreER^T2^/Nkcc1^fl/fl^/tdTomato* mice followed by transcardial perfusion 14- and 28-days post injection, n=3-5. (E) Confocal images of proliferating tdTomato^+^MCM2^+^ NSCs. Scale bar: 50µm (F) Increase of MCM2^+^ NSC clusters showing proliferating NSCs from all NSC clusters. Values represent mean ± SEM, Kruskal-Wallis-Test with posthoc Dunn’s Test, ∗p<0.05, dpi=days post injection. (G) Confocal images of NSC-NSC and NSC-N cluster. Scale bar: 10µm, Images: NSC-NSC (1-4 NSCs); NSC-N (1 NSC and 2-5 precursor). (H) Percentage increase of tdTomato^+^ NSC clones from all clusters 28 days post injection. Values represent mean ± SEM, Kruskal-Wallis-Test with posthoc Dunn’s Test, ∗p<0.05, dpi=days post injection. (I) Percentage decrease of tdTomato^+^ N clones from all clusters 28 days post injection. Values represent mean ± SEM, Kruskal-Wallis-Test with posthoc Dunn’s Test, ∗p<0.05, dpi=days post injection. (J) Percentage increase of tdTomato^+^ NSC-NSC clusters from all maintenance (M) clusters (including NSC-NSC, NSC-N, NSC-A, NSC-N-A) 28 days post injection due. Values represent mean ± SEM, Kruskal-Wallis-Test with posthoc Dunn’s Test, ∗p<0.05, dpi=days post injection. (K) Percentage decrease of tdTomato^+^ NSC-N clusters from all maintenance clusters 28 days post injection. Values represent mean ± SEM, Kruskal-Wallis-Test with posthoc Dunn’s Test, ∗p<0.05, dpi=days post injection. (L) Confocal immunofluorescence images showing a cluster with two tdTomato^+^ NSC clones. Scale bar: 10µm. Increase of all clusters containing at least two tdTomato^+^ NSCs from all tdTomato^+^ NSC clusters after *Nkcc1ko*. Values represent mean ± SEM, Kruskal-Wallis-Test with posthoc Dunn’s Test, ∗p<0.05, dpi=days post injection. (M) Confocal immunofluorescence images showing clusters with at least three tdTomato^+^ NSCs. Scale bar: 10µm. Increase of all clusters containing three tdTomato^+^ NSCs from all NSC clusters 14 days post injection after *Nkcc1ko*. Values represent mean ± SEM, Kruskal-Wallis-Test with posthoc Dunn’s Test, ∗p<0.05, dpi=days post injection. (N) Confocal immunofluorescence images showing dentate gyrus from 3m Ctrl and 3m *Nkcc1ko*. Scale bar 10µm. Experimental design for long-term changes after *Nkcc1ko*. (O) Quantification of all tdTomato^+^ NSCs in 3-month-old control and 3-month-old *Nkcc1ko* mice 6 weeks post injection. Values represent mean ± SEM, Kruskal-Wallis-Test with posthoc Dunn’s Test, ∗p<0.05. (P) Increased ratio of tdTomato^+^ NSCs to all tdTomato^+^ cells after *Nkcc1ko*. Values represent mean ± SEM, Kruskal-Wallis-Test with posthoc Dunn’s Test, ∗p<0.05. (Q) Decrease in all tdTomato^+^ neurons due to *Nkcc1ko*. Values represent mean ± SEM, Kruskal-Wallis-Test with posthoc Dunn’s Test, ∗p<0.05. (R) Decrease in the ratio of all tdTomato^+^ neurons to all tdTomato^+^ cells due to *Nkcc1ko*. Values represent mean ± SEM, Kruskal-Wallis-Test with posthoc Dunn’s Test, ∗p<0.05.

Furthermore, we found that clusters containing multiple NSCs (NSC-NSC clusters), in relation to the maintenance clusters containing clusters with NSCs (NSC-NSC: only stem cells, NSC-N: stem cells and IPCs, NSC-A: stem cells and astrocytes, NSC-N-A: stem cells, IPCs and astrocytes), predominantly emerged upon knockout of *Nkcc1*, whereas in the controls asymmetric division of NSCs with the formation of NPCs (NSC-N clusters) prevailed (Fig. 1J/K). This suggested that *Nkcc1ko* likely promoted symmetric NSC division (expansion stem cell pool), while clusters containing both NSCs and NPCs declined. Within these NSC clusters, clusters consisting of 2 or 3 NSCs were predominantly identified (Fig. 1L/M). To investigate the long-term effects of *Nkcc1ko*, mice were injected with tamoxifen for 5 days, and the number of NSCs- and neurons was analyzed 1.5 months later (Fig. 1N). The results showed that, on the long-term, the *Nkcc1ko* significantly increases the NSC population relative to all tdTomato-positive cells (NSCs, NPCs, neurons and astrocytes), while reducing the formation of new neurons compared to controls (Fig. 1 O-R).

The *Nkcc1ko* in NSCs led to a shift in the cluster formation towards symmetric NSC division. Over all, our data suggest that the chloride cotransporter NKCC1 has a significant impact on the fate of NSCs, favoring symmetric NSC division, thereby increasing the NSC pool.

### *Nkcc1* knockout reduces *Sox11* expression in NSCs

To elucidate the mechanisms involved in altered symmetric NSC division and reduction in neurogenesis *Nkcc1ko*, we performed single-cell RNA sequencing of NSCs from *Nkcc1ko* and control animals (Fig. 2A). We extracted the hippocampus and isolated individual cells from the dissociated tissue. After sequencing each tdTomato-positive cell, we assigned them to the NSC category using characteristic markers described by Shin et al. and conducted an analysis of differentially expressed genes between the two groups (Shin *et al*, 2015). The analysis revealed a significant reduction of the expression of *Sox11* in the NSCs after *Nkcc1ko* (Fig. 2B-D). Additionally, the Gene Ontology (GO) term analysis indicated that genes associated with neurogenesis were significantly reduced by the knockout, further supporting our findings that NKCC1 is a relevant contributor to the fate of the NSCs (Fig. 2E). Among the differential expressed genes, Sox11 was identified as an important regulator of neurogenesis and NSC division (Mu *et al*., 2012; Wang *et al*., 2013).

**Figure 2.**
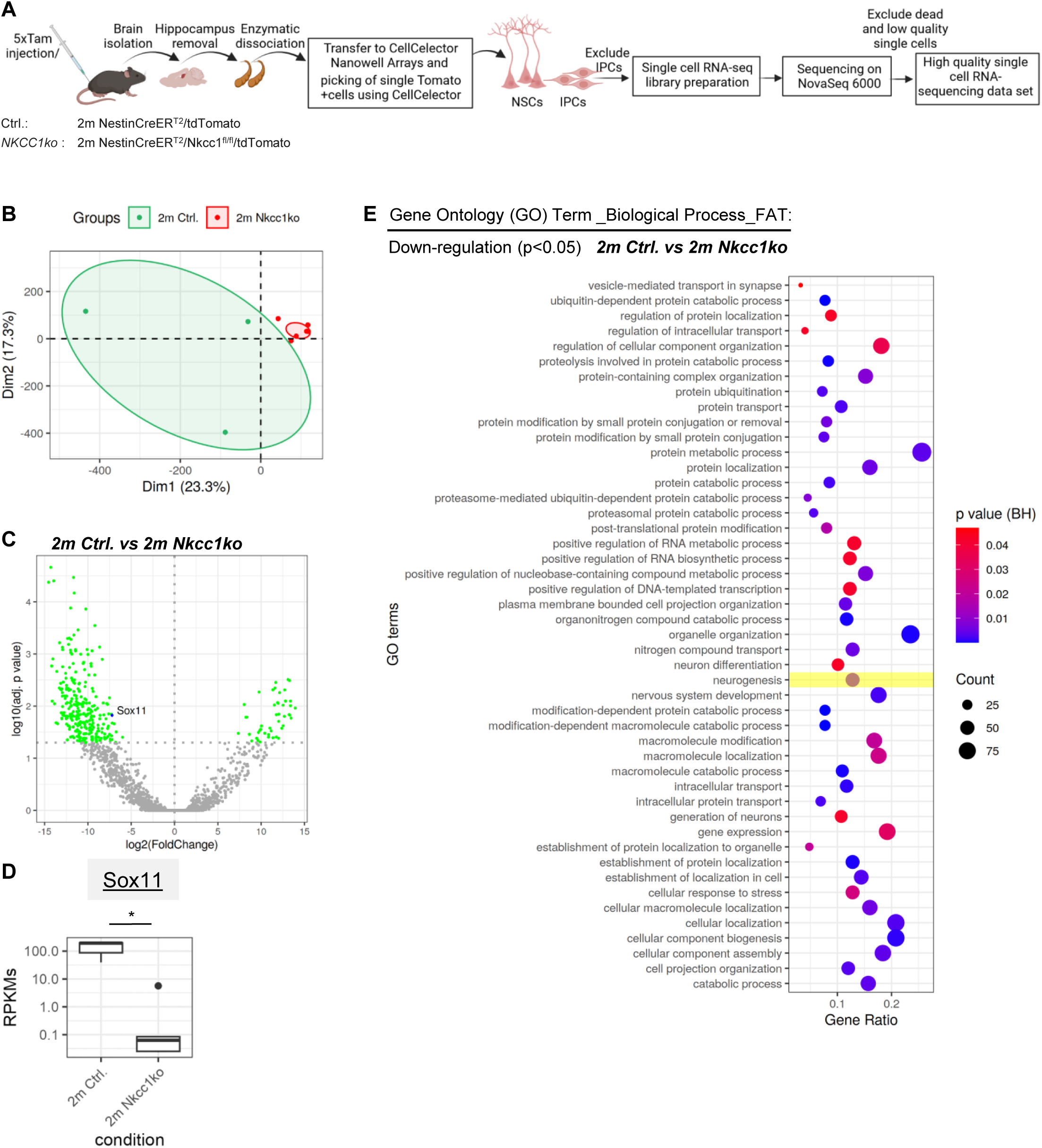
Individual RNA single cell sequencing of NSCs. (A) Schematic illustration of individual RNA single cell sequencing. (B) NSCs from 2-month-old control and *Nkcc1ko* mice cluster distinctly in a principal component analysis (PCA). (C) Down-(left side) and up-regulated (right side) genes of single NSCs derived from 2-month-old control and *Nkcc1ko* mice, highlighting down-regulated *Sox11* gene, plot shows FoldChange vs. adj. p-value. (D) Among the DEGs, the transcription factor Sox11, which is crucial for neuronal lineage of NSCs, was found to be down-regulated through *Nkcc1* knockout. (E) Top downregulated Gene Ontology (GO) terms between single NSCs of 2m control and 2m knockout mice revealed NKCC1-dependent regulation of neurogenesis. The vertical axis represents the name of the GO-terms and the horizontal axis shows the gene ratio. The size of the dots denote the fraction of affected genes in the respective GO-term while the color corresponds to the *p*-value.

### *Sox11* knockout induces NSC proliferation and symmetric NSC division

Next, we examined the possible role of Sox11 as a putative downstream factor associated with the age-dependent regulatory effects of NKCC1 on the self-renewal of NSCs. Sox11 - a member of the SoxC transcription factor family - is predominantly expressed in neurogenic areas of the adult brain and is involved in the regulation of specific gene expression programs in adult neurogenesis at the stage of immature neurons (Haslinger *et al*., 2009; Mu *et al*., 2012; Wang *et al*., 2013).

To understand the extent to which Sox11 influences the proliferation of NSCs after *Nkcc1ko*, tamoxifen was administered to *Nestin-CreER^T2^/Nkcc1^fl/fl^/tdTomato* mice for five consecutive days, followed by an analysis of proliferating Sox11-positive NSCs (Fig. 3A). Results show a significant reduction in the number of Ki67-Sox11-positive NSCs, both in absolute numbers and as a percentage of proliferating tdTomato-positive NSCs (Fig. 3B/C). Furthermore, our data suggest that in the control group, approximately 50% of proliferating NSCs showed Sox11 levels, whereas in knockout mice, Sox11 levels in proliferating NSCs was significantly reduced (Fig. 3B/C).

**Figure 3.**
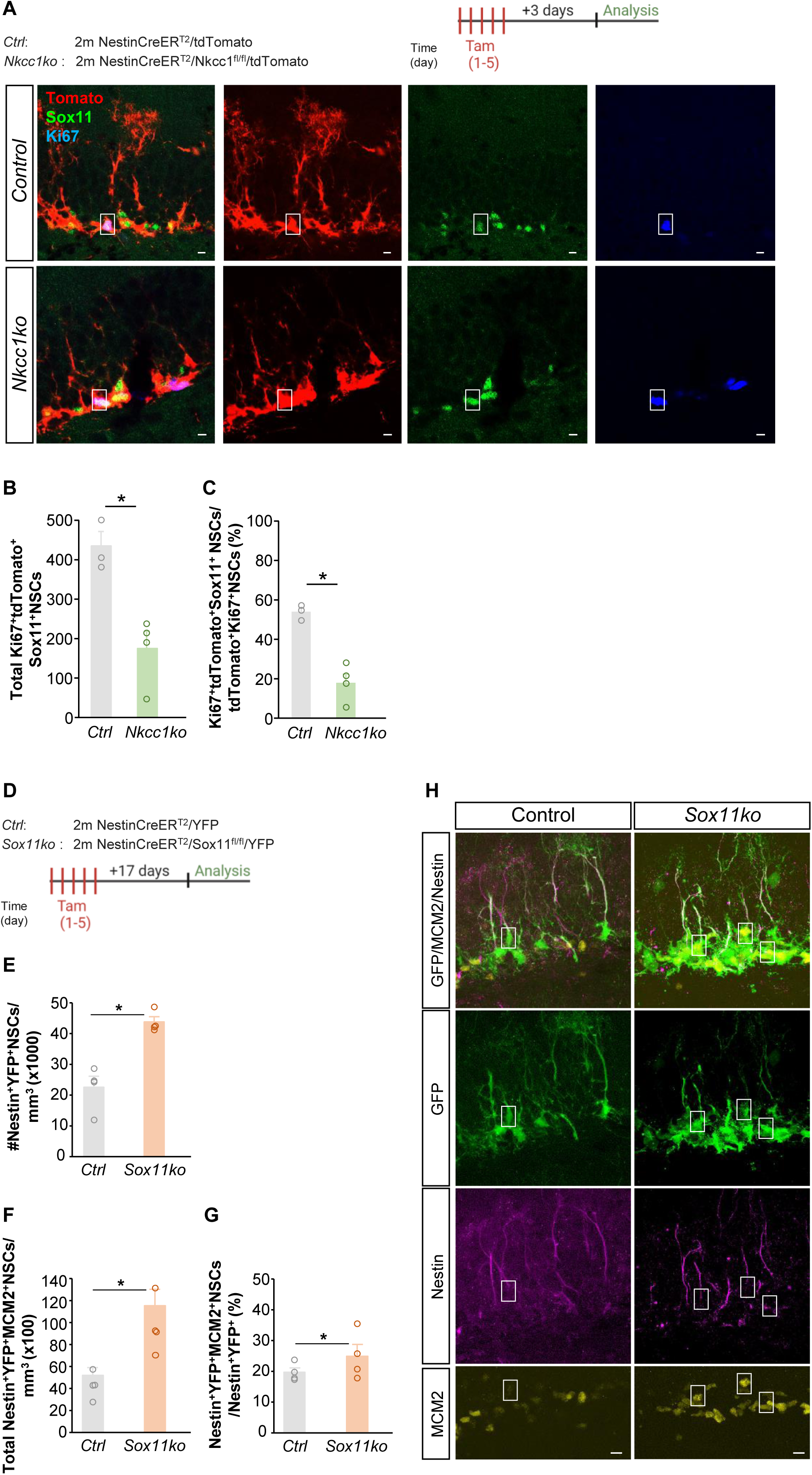
*Sox11* knockout induces NSC proliferation and self-renewal (A) Experimental design using 2-month-old *NestinCreER^T2^/tdTomato* mice and *NestinCreER^T2^/Nkcc1^fl/fl^/tdTomato* mice. Confocal immunofluorescence images showing proliferating Sox11^+^ NSCs. Scale bar: 10µm. (B) Quantification of all Ki67^+^tdtomato^+^Sox11^+^ NSCs in 2-month-old *NestinCreER^T2^/tdTomato* mice and 2-month-old *NestinCreER^T2^/Nkcc1^fl/fl^/tdTomato* mice, after tamoxifen application. Values represent mean ± SEM, two-tailed t-Test, ∗p<0.05. (C) Decreased ratio of proliferating Ki67^+^tdTomato^+^Sox11^+^ NSCs to all proliferating Ki67^+^tdTomato^+^ NSCs. Values represent mean ± SEM, two-tailed t-Test, ∗p<0.05. (D) Experimental design using 2-month-old *NestinCreER^T2^/YFP* mice and *NestinCreER^T2^/Sox11^fl/fl^/YFP* mice. (E) Quantification of all Nestin^+^YFP^+^ NSCs in 2-month-old control and *Sox11ko* (n=4) mice. Values represent mean ± SEM, Mann-Whitney U-Test, ∗p<0.05. (F) Quantification of proliferating Nestin^+^YFP^+^MCM2^+^ NSCs. Values represent mean ± SEM, Mann-Whitney U-Test, ∗p<0.05. (G) Percentage of Nestin^+^YFP^+^MCM2^+^ NSCs from all Nestin^+^YFP^+^ NSCs cells. Values represent mean ± SEM, Mann-Whitney U-Test, ∗p<0.05. (H) Confocal immunofluorescence images showing YFP-Sox2-positive NSCs after *Sox11ko*. Scale bar: 10µm.

To examine the extent to which Sox11 influences not only proliferation but also the generation of new NSCs by symmetric self-renewal, we knocked out the transcription factor Sox11 in NSCs. For this, *Nestin-CreER^T2^/Sox11^fl/fl^/YFP* mice were injected intraperitoneally with tamoxifen for five days, and transcardially perfused 17 days post-injection (Fig. 3D). Our results show that the number of NSCs significantly increases following *Sox11* knockout (*Sox11ko*) (Fig. 3E). This increase was also reflected in the proliferation of NSCs, both in absolute numbers and as a percentage relative to the NSC population (Fig. 3F/G).

These findings strongly suggest that NKCC1 and the downstream effector Sox11 are key factors determining the fate of NSCs in the adult hippocampus. In particular, their reduced level leads to symmetric division and expansion of the NSC pool.

### Age-dependent NKCC1 decline enhances symmetric stem cell division

Given the substantial decrease in neurogenesis with aging (Encinas *et al*., 2011; Kuhn *et al*, 1996; Walter *et al*, 2011) and the reported potential for symmetric NSC division during this process, we next determined whether these changes might be associated with changes in NKCC1 levels (Ibrayeva *et al*., 2021). For this, we used *Nestin-GFP* mice and analyzed the expression of the chloride cotransporters NKCC1 and KCC2 in NSCs by at 2, 8, and 20 months of age (Fig. 4A/B). Our analysis indicates that the expression of the chloride cotransporter NKCC1 significantly decreases with aging, while KCC2 remains unchanged (Fig. 4C). In order to determine whether changes in the levels of NKCC1 are directly involved in age-related changes of NSC self-renewal, we used the clonal analyses data sets previously described (see Fig. 1) and compared them with the 20-month-old *NestinCreER^T2^/tdTomato* controls (Fig. 4D). Interestingly, the 20-month-old controls showed the same trends as the 2-month-old *Nkcc1ko* animals. There was an increase in NSC clones across all clusters in the 20-month-old controls and *Nkcc1ko* mice as well as a decrease in neuronal progenitor clones compared to 2-month-old *NestinCreER^T2^/tdTomato* controls (Fig. 4E/F). NSC cluster analysis also revealed similar patterns in knockout compared to aged animals (Fig. 4G/H). This indicates that both knockout and aged NSCs exhibit a significantly increased symmetric NSC division concomitant with a decrease in asymmetric division. To further evaluate long-term changes of *Nkcc1ko*, we compared the 2-month-old *NestinCreER^T2^/tdTomato* and 2-month-old *NestinCreER^T2^/Nkcc1^fl/fl^/tdTomato* animals from Fig. 1, as previously described, and compared these datasets with 8-month-old *NestinCreER^T2^/tdTomato* controls (Fig. 4I). An increase in NSCs relative to all tdTomato-positive cells was observed at 1.5-month post injection in NSCs from both knockout and middle-aged mice (Fig. 4J/K). Additionally, a comparison of neurons between the 2-month-old controls and *Nkcc1ko* as well as 8-month controls revealed a similar decrease in both groups (Fig. 4L/M).

**Figure 4.**
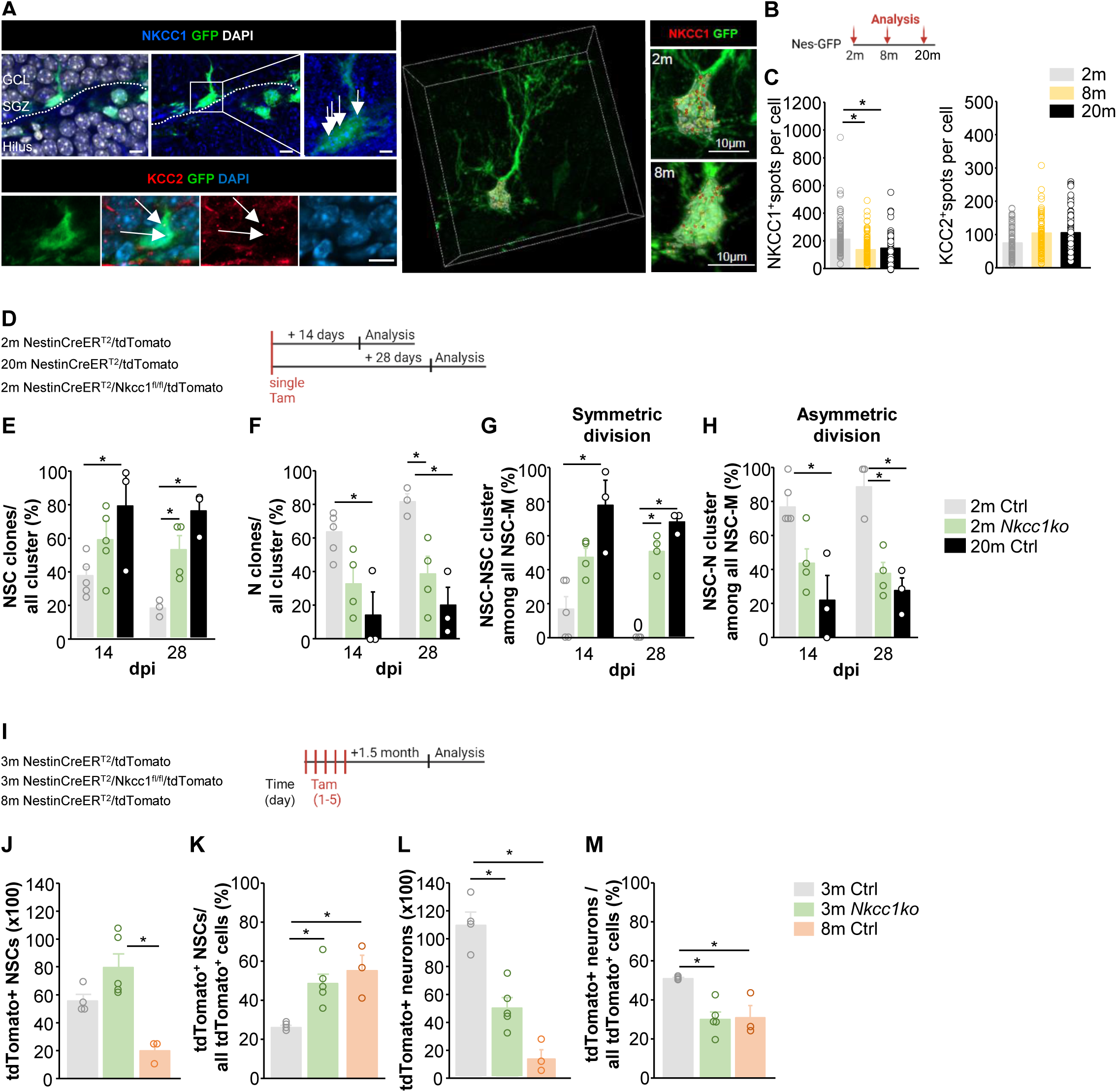
Age-dependent increase of NSC self-renewal (A) Confocal immunofluorescence images showing NKCC1 (upper panel) and KCC2 (lower panel) expression in the Nestin GFP^+^ NSCs. Scale bar: 10μm (B) Experimental design using 2-, 8- and 20-month-old *Nestin-GFP* mice. (C) Reduction of NKCC1 in NSCs in the dentate gyrus during aging but no changes of KCC2. Values represent mean ± SEM. Mice: 2 months (n=5), 8 months (n=6) and 16 months (n=5); cells: 98-125. The graph shows the mean values (± SEM), ANOVA with posthoc Tukey Test, ∗p<0.05. (D) Scheme to analyse the long-term changes 6 weeks after the last tamoxifen injection. (E) Percentage increase of tdTomato^+^ NSC clones from all clusters 28 days post injection due to aging and *Nkcc1ko*. Values represent mean ± SEM, Kruskal-Wallis- Test with posthoc Dunn’s Test, ∗p<0.05, dpi=days post injection. (F) Percentage decrease of tdTomato^+^ N clones from all clusters 28 days post injection of aging and *Nkcc1ko*. Values represent mean ± SEM, Kruskal-Wallis-Test with posthoc Dunn’s Test, ∗p<0.05, dpi=days post injection. (G) Percentage increase of tdTomato^+^ NSC-NSC clusters from all maintenance clusters 14- and 28-days post injection due to aging and *Nkcc1ko*. Values represent mean ± SEM, Kruskal-Wallis-Test with posthoc Dunn’s Test, ∗p<0.05, dpi=days post injection. (H) Percentage decrease of tdTomato^+^ NSC-N clusters from all maintenance clusters 14- and 28-days post injection due to aging and *Nkcc1ko*. Values represent mean ± SEM, Kruskal-Wallis-Test with posthoc Dunn’s Test, ∗p<0.05, dpi=days post injection. (I) Experimental design for long-term changes after *Nkcc1ko*. (J) Quantification of all tdTomato^+^ NSCs in 3- and 6-month-old control and 3-month- old *Nkcc1ko* mice. Values represent mean ± SEM, ANOVA with posthoc Tukey Test, ∗p<0.05. (K) Increase in the ratio of tdTomato^+^ NSCs to all tdTomato^+^ cells. Values represent mean ± SEM, ANOVA with posthoc Tukey Test, ∗p<0.05. (L) Decrease in all tdTomato^+^ neurons due to aging and *Nkcc1ko*. Values represent mean ± SEM, ANOVA with posthoc Tukey Test, ∗p<0.05. (M) Decrease in the ratio of all tdTomato^+^ neurons to all tdTomato^+^ cells due to aging and *Nkcc1ko*. Values represent mean ± SEM, ANOVA with posthoc Tukey Test, ∗p<0.05.

Given the similar trends observed between aging and knockout conditions, we suggest that NKCC1 reduction plays an important role in the age-related decline in neurogenesis.

## DISCUSSION

Our study indicates that the chloride cotransporter NKCC1 is a key regulator of Nestin-positive adult stem cell division. Knocking out *Nkcc1* in NSCs enhances proliferation and promotes symmetric NSC division in the adult dentate gyrus leading to an expansion of the NSC pool. We identified the transcription factor Sox11 as a direct downstream mechanism, involved in the regulation of neurogenesis. In the absence of *Nkcc1*, Sox11 levels were reduced in proliferating NSCs. Furthermore, the knockout of *Sox11* in NSCs confirmed its role in increasing both proliferation and the NSC pool.

Various studies (Bao *et al*., 2017; Song *et al*., 2012; Zhang *et al*., 2023) have already demonstrated the role of the neurotransmitter GABA and the GABA_A_ receptor in the proliferation and mode of division of NSCs. In this context, it was shown that a decrease in GABA, the loss of function of the GABA_A_ receptor due to the depletion of its delta subunit (Song *et al*., 2012), as well as *Nkcc1* knockout in HopX-positive adult stem cells (Zhang *et al*., 2023), result in increased stem cell proliferation. Consistent with these findings, our study demonstrated an increase in proliferation due to the *Nkcc1ko* in NSCs. Clearly, this effect is predominant across various stem cell populations. In contrast to the study by Zhang et al. focusing on the HopX stem cells (Zhang *et al*., 2023), our findings indicate that, following *Nkcc1ko*, NSCs undergo symmetric NSC division to expand the NSC pool. This leads to impaired hippocampal neurogenesis. Their study demonstrated an increase in NPCs, which consequently contributes to enhanced neurogenesis. Stem cells are a heterogeneous group of cells (Bottes *et al*, 2021; Obernier & Alvarez-Buylla, 2019; Pilz *et al*., 2018), and this heterogeneity might account for the differences observed, as previously described by (Ibrayeva *et al*., 2021), when comparing Ascl1-positive and Nestin-positive stem cells. Ascl1-positive stem cells are described as neural stem/progenitor cells (NSPCs), meaning that these cells are already in an intermediate state between stem and progenitor cells.

The NSCs are mainly in a dormant state. The mechanisms that lead to NSC activation and additionally to symmetric division are still poorly understood. In our study, we observed that the enhanced symmetric NSC division was associated with a decrease in the transcription factor Sox11. This association was confirmed by the *Sox11* knockout, which resulted in an increase in the NSC pool. Previous studies have already demonstrated the importance of *Sox11* in stem- and progenitor cells (Haslinger et al., 2009; Mu et al., 2012). Our study shows that a high proportion of proliferating NSCs express Sox11.

After the *Nkcc1ko*, NSCs showed a reduction in the number of Sox11-level NSCs within the proliferating population. This suggests that Sox11 levels may play a crucial role in determining NSC fate, whether by promoting the generation of new neurons or by expanding the NSC pool.

The NSC pool in the hippocampal dentate gyrus significantly decreases during aging, leading to a reduction in the formation of new neurons. This decline in the NSC pool is particularly pronounced during the transition from young to middle age (Encinas *et al*., 2011; Ibrayeva *et al*., 2021; Wu *et al*., 2023). Subsequently, the absolute number of NSCs steadily decreases. This decline is partly attributed to the decrease in NSC proliferation (Encinas *et al*., 2011; Heine *et al*, 2004; Kuhn *et al*., 1996; Walter *et al*., 2011). Additionally, studies (Bonaguidi *et al*., 2011; Obernier *et al*, 2018) have shown that, apart from the age-related changes in proliferation, the mode of NSC division occur both symmetrically to maintain the NSC pool and asymmetrically to promote the formation of new neurons. In this context Ibrayeva et al. showed that Nestin-positive long-term stem cells increased symmetric division with increasing age, while the short-term Ascl1-NSPCs exhibit low self-renewal capacity (Ibrayeva *et al*., 2021). Pilz et al. demonstrated that NSPCs undergo limited rounds of symmetric and asymmetric cell divisions, which promote neurogenesis, after which they are lost (Pilz *et al*., 2018). The mechanisms underlying changes in modes of cell division are not yet fully understood. We found that the chloride cotransporter NKCC1 is expressed in murine hippocampal NSCs and decreases significantly during aging, while its counterpart KCC2 remains stable. When comparing young *Nkcc1ko* NSCs to aged NSCs, we observed strong parallels, particularly regarding the increase in symmetric NSC divisions and decrease in neurogenesis. These findings suggest that NKCC1 expression plays a key role in preserving the NSC pool during aging.

Our study reveals a novel regulatory pathway of symmetric NSC division, significantly contributing to the understanding the age-related decline in the NSC pool and in adult neurogenesis.

## Limitations of the study

Our study provides evidence for the role of NKCC1 and its downstream effector Sox11 in the fate and proliferative capacity of NSCs in the adult hippocampus, however, the precise mechanisms underlying the effects of *Nkcc1ko* and the associated downregulation of *Sox11* in NSCs remain to be fully elucidated. In particular, it is still unclear whether the observed reduction in *Sox11* expression is primarily driven by altered intracellular chloride concentrations, the loss of depolarizing GABAergic signals, or a combination of both. Since NKCC1 plays a key role in maintaining the chloride gradient necessary for GABA-induced depolarization, its deletion could disrupt critical activity-dependent signaling pathways that regulate transcription factor expression and NSC fate decisions. At present, the molecular cascade linking these physiological changes to transcriptional regulation remains unclear. Therefore, future investigations will be essential to identify the specific signaling pathways and gene regulatory networks regulated by NKCC1.

## ACKNOWLEDGMENTS

S.K. received support from Deutsche Forschungsgemeinschaft (DFG; grants KE 1914/2-1; Interdisciplinary Center for Clinical Research Jena (Woman in science FF06); D.C.L. received support from the Deutsche Forschungsgemeinschaft (LI 858/9-2). S.A. is a member of the research training group 2162 “Neurodevelopment and Vulnerability of the Central Nervous System” funded by the Deutsche Forschungsgemeinschaft (270949263/ DFG GRK2162/2). K.K. received support from Deutsche Forschungsgemeinschaft (KI 1816/6-2 #442107075). A.B., M.W. were supported by funding from the Foundation “Else Kröner-Fresenius-Stiftung” within the Else Kröner Graduate School for Medical Students “Jena School for Ageing Medicine; J.B., L.K.; M.R. were supported by the Interdisciplinary Center for Clinical Research Jena. The Core Facility Next Generation Sequencing (Ivonne Goerlich) of the FLI is gratefully acknowledged for their technological support in library preparation and sequencing. Created in BioRender. Keiner, S. (2025) https://BioRender.com/lc9n1f2.

## AUTHOR CONTRIBUTIONS

Conceptualization, S.K.; data collection and methodology, A.B., S.A., M.R., J.B., L.K., M.G., M.H., M.W., G.S., J.M., AL.F., T.F.; data analysis and bioinformatics, M.G., S.K., supervision, S.K.,;writing – review & editing, I.S., K.K, H.H., E.S., CA.H., K.H., C.S., DC.L., S.K. project coordination, S.K. All authors commented on the manuscript.

## STAR ★ Methods

Key resources table

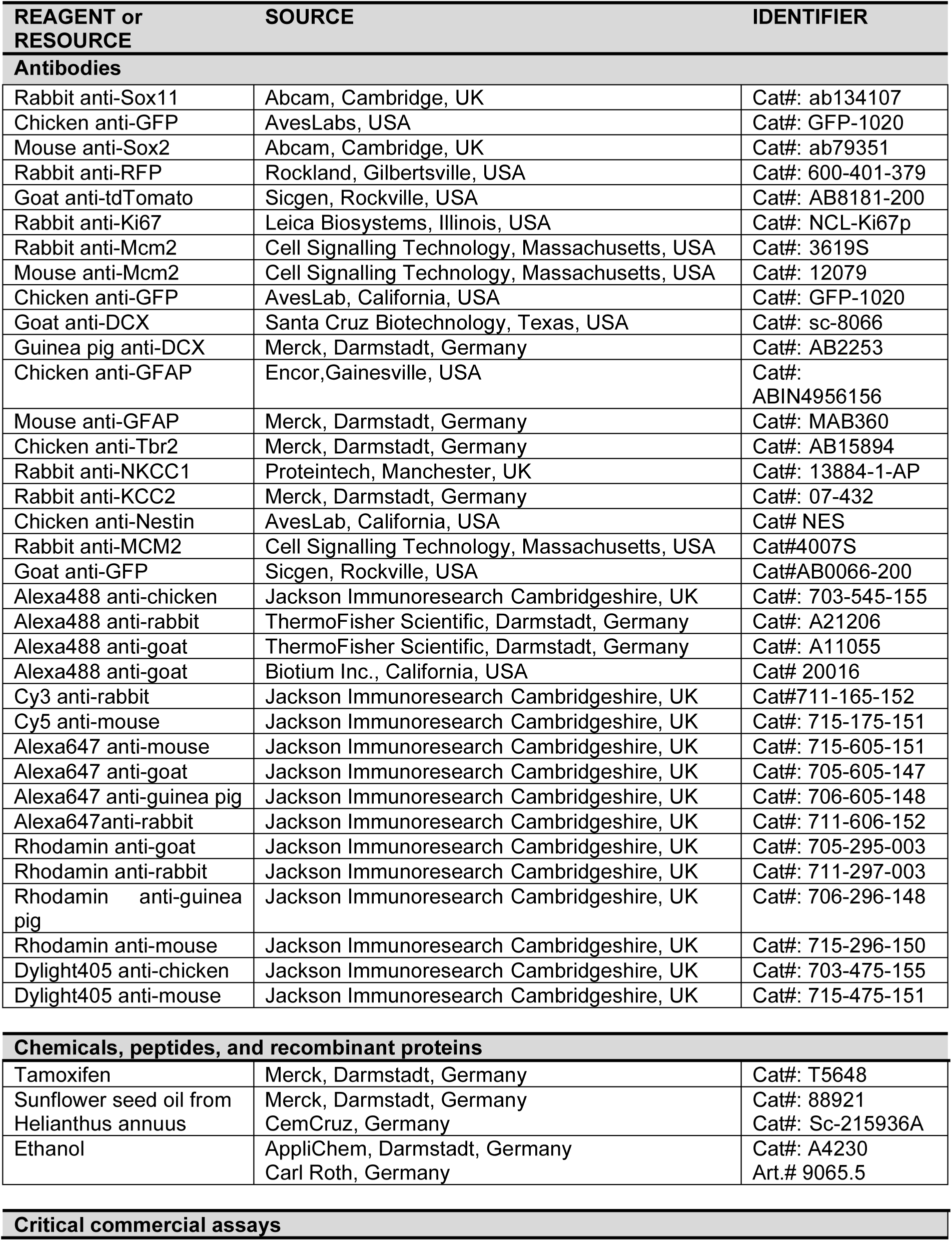
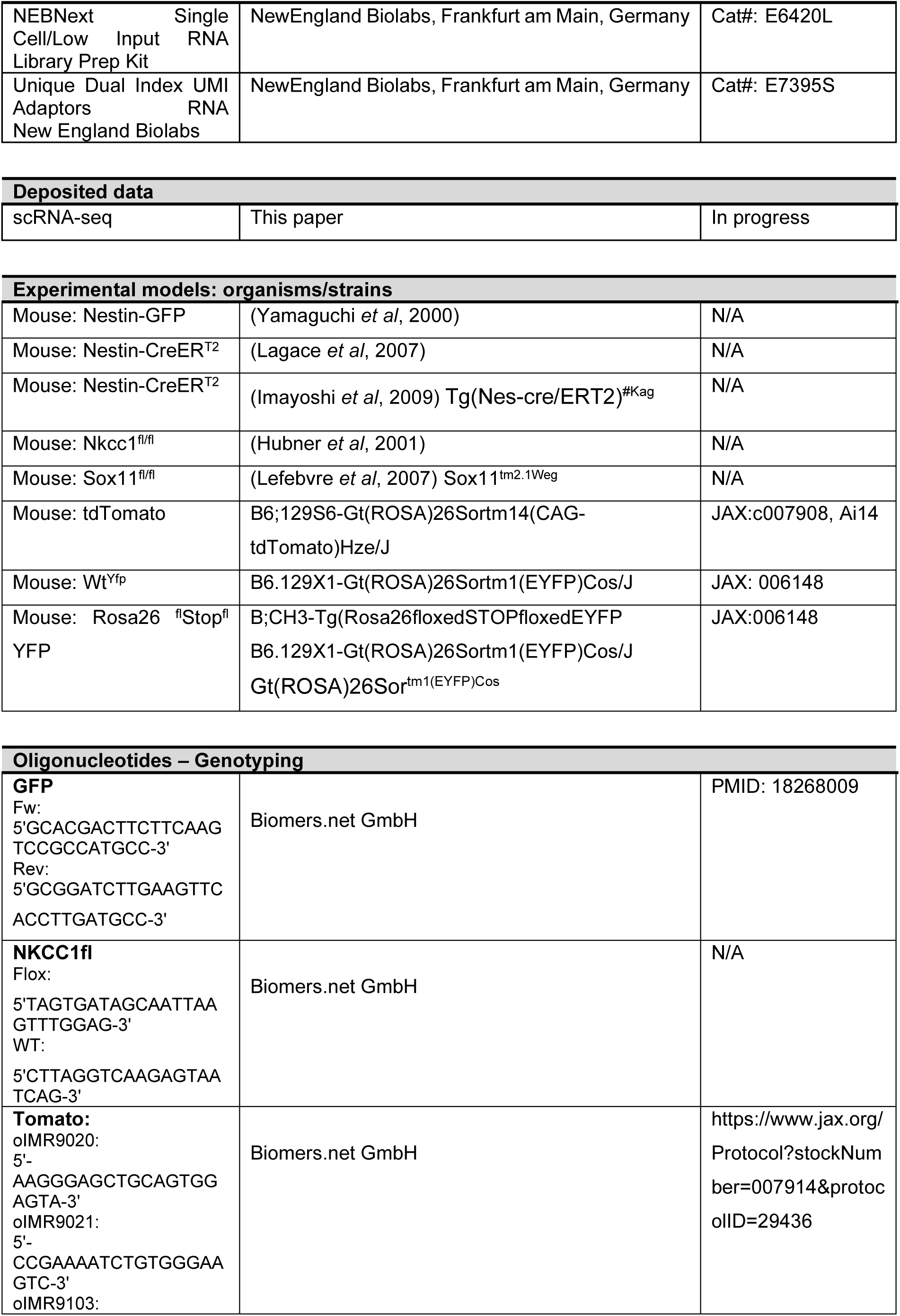
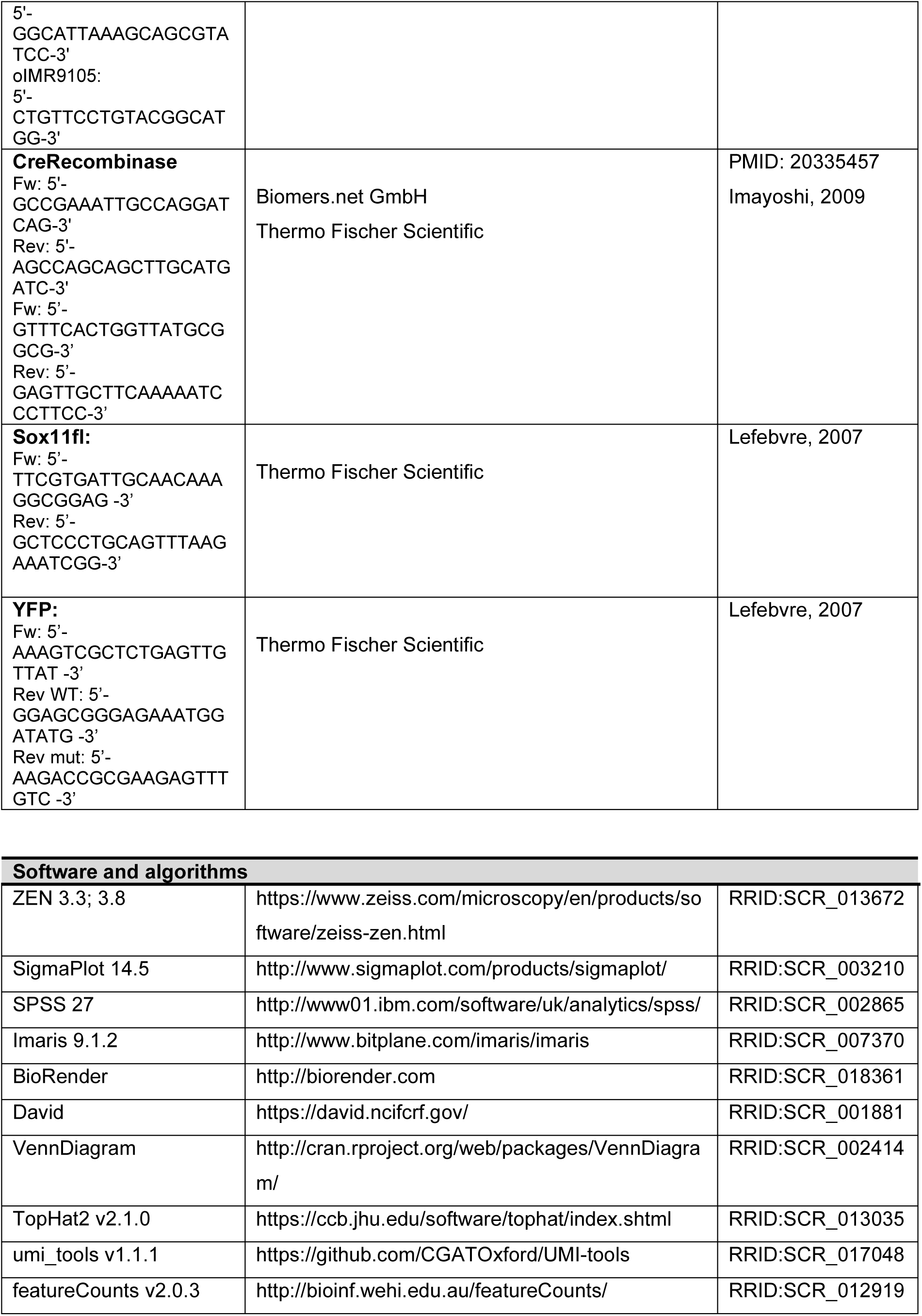
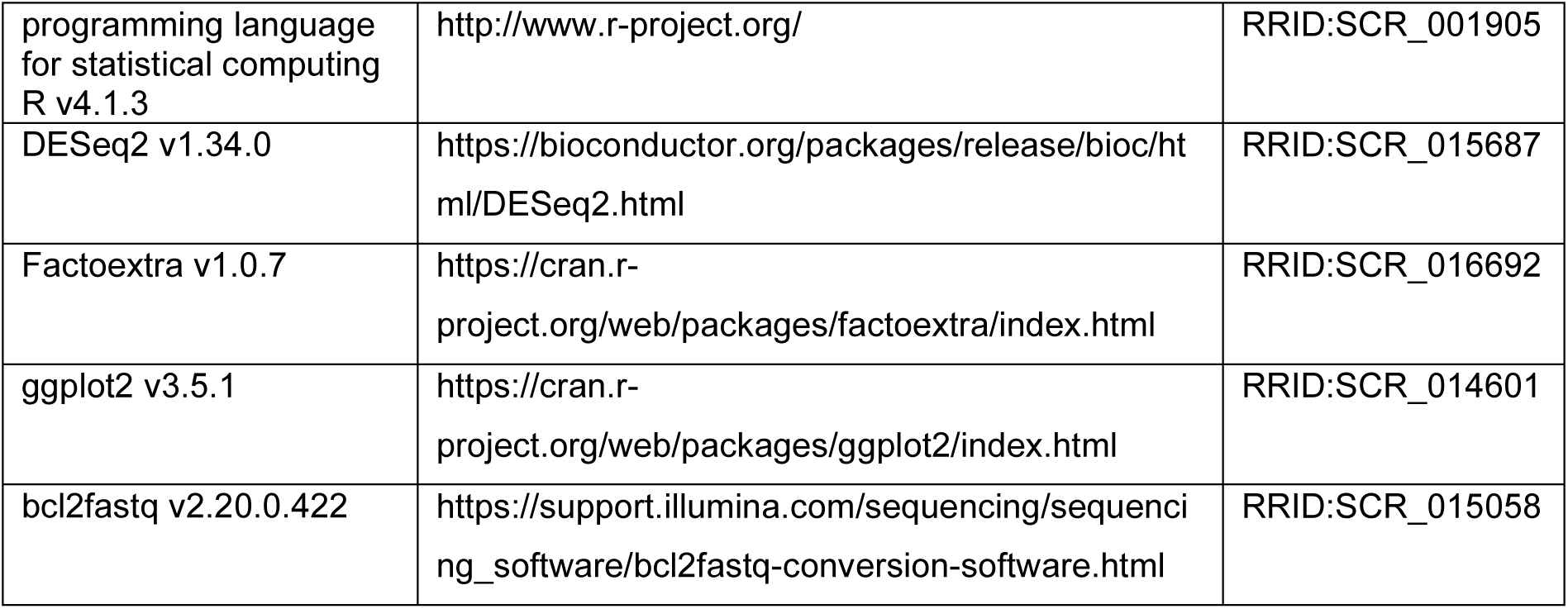

## RESOURCE AVAILABILITY

### Lead contact

Further information including requests for resources and reagents should be directed to the lead contact, Silke Keiner.

## Materials availability

The study did not generate new reagents.

## EXPERIMENTAL MODEL AND SUBJECT DETAILS

### Animals and tamoxifen administration

All animal procedures were performed in accordance with the current European guidelines and approved by the ethics committee of the local governments. Mice were housed in groups of 2-5 in standard cages (54cm/38cm/19cm) under controlled laboratory conditions (22-24°C temperature, 30-60% humidity), and in a 14h light/10h dark cycle with free access to food and water. All mice used in the study were backcrossed to the C57BL/6 background to ensure the reproducibility of clonal induction with specific doses of tamoxifen.

The *Nestin-GFP* mice (Yamaguchi *et al*., 2000) were used for the analysis of age-associated changes in NSCs under physiological conditions.

Clonal analysis of NSCs was performed using *NestinCreER^T2^* mice provided by Prof. A. Eisch, Department of Psychiatry, University of Texas Southwestern Medical Center, Dallas, USA (Lagace *et al*., 2007). The genetically modified *Nkcc1*^flox^ mouse line was kindly provided by Prof. CA. Hübner (Institute of Human Genetics, University Hospital Jena, Germany, (Hubner *et al*., 2001) on a C57BI/6J background.

The *Nestin-CreER^T2^* and *Nestin-CreER^T2^/Nkcc1^fl/fl^* mice were crossbred with fluorescent reporter mice (Strain: B6;129S6-Gt(ROSA)^26Sortm14(CAG-tdTomato)Hze^/J Stock No: 007908, Ai14) for clonal analysis and quantification of NSCs, precursor cells and neurons. The *Nestin-CreER^T2^/Sox11^fl/fl^* mice were crossbred with fluorescent reporter mice (Strain: B6.129X1-Gt(ROSA)26Sortm1(EYFP)Cos/J) for quantification of NSC populations.

Animals from different litters were checked for the reporter/driver combination to ensure there was no recombination in the adult SGZ in the absence of tamoxifen. For the clonal analysis, a stock of tamoxifen (15mg/ml) was prepared in a 1:9 ratio of ethanol to sunflower oil at 37°C with vortexing. A single dose of the tamoxifen solution (60mg/kg body weight) was injected intraperitoneally into the mice. Injected animals showed no signs of distress.

To verify the total number of tdTomato^+^ NSCs and neurons, a single daily dose of tamoxifen (30mg/ml) was injected over the first 5 days at a concentration of 180mg/kg body weight.

For generating *Sox11ko* mice, *Nestin-CreER^T2^* (Imayoshi *et al*, 2006) were crossbred with B;CH3-Tg(Rosa26floxedSTOPfloxedEYFP) (Potzner et al., and B6;CH3-*Sox11^flox/flox^* (Penzo-Mendez et al., 2007). Tamoxifen solution (10mg/ml) was prepared in ethanol and Sunflower oil (1:9). To induce the deletion, mice were injected intraperitoneally (30mg/kg body weight) 2 times a day for 5 days.

For genotyping, the DNA was isolated from ear notches in a 25mM NaOH, 0.2mM EDTA solution for 30min. at 95°C. After the heating incubation step the DNA was neutralized by adding 40mM Tris-HCL. Genotyping was performed with specific protocols on a Biometra Thermocycler TRIO. For detection of cre-recombinase, the PCR protocol was the following: 95°C 30sec, 61°C 30sec, 72°C 60sec (35cycles); *Nkcc1*: 94°C 30sec, 57°C 30sec, 72°C 45sec (35cycles), tdTomato: 94°C 20sec, 61°C 30sec, 72°C 30sec (35cycles).

For genotyping the DNA was isolated from ear clips lysed in 50mM Tris-HCl, 100mM EDTA and 0.5% SDS at 55°C for 1hr. The DNA was precipitated by adding isopropanol and ethanol. The pelleted DNA was dried and reconstituted in milliQ water. Genotyping was carried out using following protocols in Eppendorf mastercycler epgradient S. For NestinCre: 95°C 30sec, 53°C 30sec, 72°C 60sec (40cycles); YFP: 95°C 30sec, 58°C 30 sec, 72°C 60sec (30cycles); *Sox11* 95°C 30sec, 58°C 30sec, 72°C 60sec (40cycles).

### Immunostaining, confocal imaging and processing

Mice were killed by overdose of isoflurane and perfused transcardially with PBS and 4% paraformaldehyde. Brains were removed and post-fixed in 4% paraformaldehyde for 24h at 4°C. Afterwards brains were transferred to a 10% sucrose for 24h followed by 30% until they sunk. Brains were sectioned into 40µm sequential coronal sections. Sections were stored at -20°C in cryoprotect solution. The hippocampal sections underwent immunohistology procedures.

Immunofluorescence staining for the analysis of age-dependent changes in Nestin-GFP stem cells and proliferation was performed using the *Nestin-GFP*-mouse model. For this, every 24^th^ section was selected and sections were washed six times for 10min in TBS at room temperature. Sections were then blocked with TBS+ (TBS, 0.1% Triton-X-100 and 3% normal donkey serum) for one hour at room temperature. After blocking, sections were incubated in TBS+ and the corresponding primary antibodies for 24h at 4°C. For population analysis, primary antibodies chicken anti-GFP (1:200), guinea-pig anti-DCX (1:250), goat anti-DCX (1:50), mouse anti-GFAP (1:500) and rabbit anti-Ki67 (1:500) were used. Thereafter, brain sections were washed three times for 10min and were blocked again with TBS+ for 30min. Sections were then incubated with secondary antibodies Alexa488 anti-chicken (1:500), Alexa647 anti-rabbit (1:500), Alexa647 anti-guinea pig (1:500), Alexa647 anti-goat (1:500), Dylight anti-mouse (1:500) and Rhodamine anti-mouse (1:500), for 2h at room temperature and in the dark.

Sections were washed three times in TBS and mounted on slides using gelatine. Cell nuclei were stained by immersing the slides in a DAPI solution for 3-5min at room temperature in the dark. Slides were washed twice in PBS for 5min and sections were covered with Mowiol. To reduce age-related lipofuscin autofluorescence, the tissue sections of the 40- and 70-week-old animals were treated with copper II sulphate/sodium acetate.

Depending on the mouse model and the tamoxifen regime used, hippocampal sections were selected for clonal or for population analysis. For clonal analysis, all hippocampal sections were stained using immunofluorescence (Ibrayeva *et al*., 2021). For population analysis, defined sections of the hippocampus were used (Woitke *et al*, 2023).

For the analysis of the control and *Nkcc1ko* hippocampal slices, every 24^th^ section was selected and sections were washed in TBS, followed by incubation in blocking solution consisting of 0.1% Triton-X-100 and 5% normal donkey serum (NDS) in TBS (TBS+) for 1h at room temperature. Sections were then incubated in primary antibodies over night at 4°C. Following primary antibodies were used for population analysis: goat anti-tdTomato (1:300), goat anti-DCX (1:50), goat anti-GFAP (1:500), rabbit anti-RFP (1:250), mouse anti-Sox2 (1:100), chicken anti-Tbr2 (1:200), rabbit anti-Tbr2 (1:100), and rabbit anti-tdTomato (1:200). Following primary antibodies were used for clonal analysis were used following primary antibodies: goat anti-DCX (1:50), guinea pig anti-DCX (1:100), chicken anti-GFAP (1:500), rabbit anti-Mcm2 (1:500), mouse anti-Mcm2 (1:500), rabbit anti-RFP (1:250), mouse anti-Sox2 (1:100), chicken anti-Tbr2 (1:150), goat anti-tdTomato (1:300). After three washing steps with TBS for 10 min each, a further incubation was performed first in TBS+ only for 30 min and then in TBS+ containing secondary antibodies for 2h. Sections were washed 3 times for 10min in TBS. Slices of 8-month-old mice were additionally incubated in distilled water for 10min, followed by a 10min incubation in copper II sulphate/sodium acetate. Sections were then washed three times before embedding. After staining, hippocampal sections were mounted. For DAPI staining, the slides were incubated in DAPI solution for 5min in the dark. Afterwards, slices were washed twice in TBS for 5min each. The microscope slides were covered using Mowiol.

Quantification of the tdTomato^+^ cells after *Nkcc1ko* was performed using every 24^th^ section throughout the entire dentate gyrus. For clonal analysis, every section was processed. TdTomato^+^ cells were identified using confocal laser scanning microscopy (LSM710 / LSM900, Carl Zeiss AG, Jena, Germany) and acquired as a z-stack on a Zeiss confocal system under 40x magnification (stitching was done at 0.5 overlap).

To analyse changes in NSCs after *Sox11* knockout in adult mice (8- to 9-weeks-old), animals were injected with tamoxifen twice a day for 5 consecutive days. Animals were sacrificed by CO_2_ 17 days after the last tamoxifen injection, flushed with 50ml of 1xPBS and fixed using 100ml of 4%PFA. Brains were further fixed with 4%PFA overnight, and then dehydrated in 30% sucrose solution. For sectioning, a microtome was used to cut 40μm thick coronal sections. Sections were stored at -20°C in cryoprotect solution. For staining, three hippocampal regions were selected (rostral, medial and caudal). Every 12^th^ section was picked and washed three times for 10min in 1xPBS at RT, followed by blocking with 3% donkey serum with 0.25% TritonX-100 for 1h at RT. The primary antibody solution was made of blocking solution and contained primary antibodies chicken anti-Nestin (1:500), rabbit anti-MCM2 (1:500) and goat anti-GFP (1:500). Sections were incubated for 5d at 4°C. After 5d sections were washed three times for 10min in 1xPBS and then blocked again for 1h at RT. Sections were transferred to secondary antibody solution containing Alexa488 donkey anti-goat (1:1000), Cy3 donkey anti-rabbit (1:1000) and Alexa647 donkey anti-chicken for 2 d. Sections were then washed and stained with DAPI and mounted with aqua polymount mounting medium. Two DGs per animal (Ctrl. n=4 and *Sox11ko* n=4) were imaged using a Zeiss Apotome microscope with a 20x objective. The RGLs were identified using Nestin staining. Recombined or activated RGLs were identified by their radial orientation and by GFP and MCM2 staining respectively. The total cell numbers were normalized to the volume of the respective DG.

### Immunofluorescence staining for NKCC1 and KCC2

Hippocampal sections were washed with TBS 6 times for 10min each. The blocking step was performed with TBS++ (0.2% Triton X-100 and 10% NDS in TBS) for 1h. The incubation with primary antibodies was performed using chicken anti-GFP (1:200), rabbit anti-KCC2 (1:100) or rabbit anti-NKCC1 (1:50) for 48h at 4°C. This was followed by 3 rinses with TBS for 10min each, blocking in TBS++ for 30min and incubation with secondary antibodies Alexa488 donkey anti-chicken (1:250) and Alexa647 donkey anti-rabbit (1:250) overnight at 4°C. All slices were further incubated in distilled water for 10min, followed by a 10min incubation in copper II sulphate/sodium acetate. Remaining steps were performed as previously described.

### Fluorescence quantification of NKCC1 and KCC2 signals

All GFP^+^ NSCs were scanned with a confocal laser scanning microscope (LSM 900, Carl Zeiss AG, Jena, Germany) with 63x objectives in Airy scan mode. The somatic part of the NSC was pre-defined as the first region of interest (ROI) and selected by cropping the images based on the nuclear signal of the NSC. The fluorescence staining was evaluated using the microscopy image analysis software Imaris (Vesion 9.1.2).

Therefore, each ROI was imported and a background subtraction was performed. The absolute intensity of the GFP signal was used to reconstruct the surface of the NSC. No automatic smoothing was used for the 3D reconstruction. The numbers of voxels were set according to the automatic threshold (excluding small, punctual fluorescence intensities). Based on the accumulated Cy5 grey scale intensities (KCC2 or NKCC1 signal), spots were defined as representing true positive spatial distributed fluorescence signals (each Alexa647 accumulation was defined as exactly one spot). The XY diameter of a spot was set to 150nm, and no PSF-elongation along the Z-axis and no further background subtraction were performed. The remaining spots were restricted to those carrying a median fluorescence intensity of Cy5 greater than 1000 (NKCC1) or 1300 (KCC2), representing a true positive fluorescence signal. The fluorescence thresholds were pre-defined based on the negative control. Morphological analysis and fluorescence quantification of NKCC1 and KCC2 was performed using Imaris.

### Clonal analysis

The clonal analysis included the volume of the entire dentate gyrus (stratum granulosum, subgranular zone). All hippocampal sections were first screened for tdTomato^+^ clones with a fluorescent microscope (LSM 900/ LSM980, Carl Zeiss AG, Jena, Germany) at 20x magnification. Clusters were defined according to previous publications Bonaguidi *et al*. 2011 and Ibrayeva *et al*. 2021 as follows: 1. Cluster containing at least one NSC (NSC-only), more than one NSC (NSC-NSC), NSCs and precursor cells (NSC-N), NSCs and astrocytes (NSC-A), NSCs and astrocytes and precursors (NSC-N-A) which are in close spatial proximity to a NSC; 2. Depleted clusters containing no NSC and only astrocytes or neurons. The maintenance cluster include NSC-NSC, NSC-N, NSC-A and NSC-N-A clusters. Approximately 8-35 clusters per dentate gyrus were evaluated, each cluster consisting of approximately 2.5 clones. Clusters were randomly distributed throughout the entire dentate gyrus. All sections with clones were then scanned with a confocal laser scanning microscope (LSM 900/ LSM980, Carl Zeiss AG, Jena, Germany) with 40x or 63x objectives. The detected clones were phenotyped according to morphological and immunofluorescence criteria, as previously described (Bonaguidi *et al*., 2011; Ibrayeva *et al*., 2021). Distance measurements were performed using the “Maximum Intensity Projection” mode of ZEN lite software. In clusters with several NSCs, the clone center is defined by the location of the most centrally situated NSC. In clusters containing no NSCs (depleted clusters), the clone center was defined by the central astrocyte or neural lineage cell. At 28 dpi, a radius of 150 μm to the clone center was set as the maximum distance. Clones were randomly distributed throughout the dentate gyrus.

### Isolation of individual cells from adult dentate gyrus

Control and *Nkcc1ko* mice, aged 2 and 6 months, were used for single-cell sequencing. Mice were killed by cervical dislocation, and brains were removed and immediately transferred to ice-cold solution A. The dentate gyrus was microdissected under stereomicroscope and transferred into 37°C dissociation solution, where it was incubated at 37°C for a maximum of 25min. During this time, the tissue was gently pipetted up and down using a 1ml pipette to loosen. Ice-cold solution B was then added in equal parts to deactivate the enzymes, and the solution was passed through a 70µm filter (Kuriakose & Xiao, 2024). The solution was centrifuged, the supernatant discarded, and cell medium was added to the cells. The cell suspension was then diluted and transferred to well plates for cell selection using a CellCelector. Only tdTomato^+^ cells were selected using the CellCelector, and only if they were free of tissue debris. Individually picked cells were immediately transferred into Lysis buffer according to manufactures instructions (NEBNext Single Cell/Low Input RNA Library Prep Kit).

For RNA analysis, the housekeeping markers Actb, Ubb, and Gapdh were initially used. To identify the NSCs, the markers *Id4, HopX, GFAP, Nestin, Apoe, Fabp* and *Aldoc* were evaluated. To exclude precursors, the markers *DCX* and *Eomes* (Tbr2) were assessed.

### Library preparation and sequencing of single cells

Sequencing of RNA samples was performed using Illumina’s next-generation sequencing methodology (Bentley *et al*, 2008). Library preparation was done with a NEBNext Single Cell/Low Input RNA Library Prep Kit in combination with NEBNext Multiplex Oligos for Illumina (Unique Dual Index UMI Adaptors RNA), following manufacturer’s instructions (New England Biolabs). Thereafter, cDNA amplification, quantification and quality check were performed using a 2100 Bioanalyzer and a High Sensitivity DNA assay (Agilent Technologies). 0.11 - 3.22 ng of amplified cDNA were used for fragmentation/end preparation step. Quantification and quality check of amplified libraries was done using a 4200 Tapestation and a D5000 assay (Agilent Technologies). Libraries were pooled and sequenced in two NovaSeq 6000 runs (1xSP + 1xS1). System runs with reagents v1.5 in 101 cycle/single-end/standard loading workflow mode. Using 19cycles in index 1 read, the index as well as the UMI sequences were obtained. Sequence information was converted to FASTQ format using the software bcl2fastq v2.20.0.422.

Mapping of reads to the reference was done with TopHat v2.1.0 (Kim *et al*, 2013) using parameters --no-convert-bam --no-coverage-search -x 1 -g 1. The human Ensembl genome version GRCh38 with annotation release 92 was used as reference (Zerbino *et al*, 2018). PCR duplicons were removed after mapping of reads based on UMIs (unique molecular identifier) using umi_tools v1.1.1 (Smith *et al*, 2017). Remaining reads per gene were counted for each sample with featureCounts v2.0.3 (Liao *et al*, 2014) using the parameter -s 0. Gene counts were further processed using the programming language R (v4.1.3). In order to find differentially expressed genes (DEG), the statistical packageDESeq2 (v1.34.0) (Love *et al*, 2014) was applied. P-values (Wald significance test) were adjusted for multiple testing according to Benjamini-Hochberg correction (FDR). Genes with adjusted p-value < 0.05 were regarded to be differentially expressed.

Principle component analysis was done using prcomp and data were visualized using fviz_pca_ind (factoextra package: 10.32614/CRAN.package.factoextra). The relation of fold-change and p-values are visualized in a volcano plot using ggplot (ggplot2 package: 10.32614/CRAN.package.ggplot2). Expression values for *Sox11* are visualized in a boxplot using ggplot.

Functional annotations of the differentially expressed genes were performed with DAVID (Database for Annotation, Visualization and Integrated Discovery)(Huang da *et al*, 2009), and results for the Gene Ontology (GO) term, biological process (GOTERM_BP_FAT) were exported. The statistical significance of the enrichment and exclusivity were assessed by two-sided Fisher’s exact test followed by the Benjamini-Hochberg adjustment (padj<0.05).

Significant GO terms (including no. of affected genes and adjusted p value) were visualized using ggplot in the programming language for statistical computing R.

## QUANTIFICATION AND STATISTICAL ANALYSIS

Statistical analyses were performed as indicated in each figure legends with the corresponding significance level and relevant statistical parameters. The IBM SPSS Software (Version 27, IBM, New York, USA) was used for statistical analyses. If data were normally distributed (Shapiro-Wilk test), then two-tailed unpaired Student’s t test (two groups) or one-way analysis of variance (ANOVA) (more than two groups) with Tukey’s multiple comparisons test were used. If data were not normally distributed, Mann-Whitney test (two groups) or Kruskal-Wallis ANOVA (more than two groups) with Dunn’s method for multiple comparisons test were used. The *Sox11* data statistics were calculated after averaging the two DGs for all animals using the Mann-Whitney test. The statistical tests employed for the single-cell RNA-sequencing analysis are integrated in the package DEseq2 (Wald significance test).

## REFERENCES

1. Abbott LC, Nigussie F (2020) Adult neurogenesis in the mammalian dentate gyrus. Anat Histol Embryol 49: 3–16

2. Bao H, Asrican B, Li W, Gu B, Wen Z, Lim SA, Haniff I, Ramakrishnan C, Deisseroth K, Philpot B, Song J (2017) Long-Range GABAergic Inputs Regulate Neural Stem Cell Quiescence and Control Adult Hippocampal Neurogenesis. Cell Stem Cell 21: 604–617 e605

3. Bentley DR, Balasubramanian S, Swerdlow HP, Smith GP, Milton J, Brown CG, Hall KP, Evers DJ, Barnes CL, Bignell HR et al (2008) Accurate whole human genome sequencing using reversible terminator chemistry. Nature 456: 53–59

4. Bonaguidi MA, Wheeler MA, Shapiro JS, Stadel RP, Sun GJ, Ming GL, Song H (2011) In vivo clonal analysis reveals self-renewing and multipotent adult neural stem cell characteristics. Cell 145: 1142–1155

5. Bottes S, Jaeger BN, Pilz GA, Jorg DJ, Cole JD, Kruse M, Harris L, Korobeynyk VI, Mallona I, Helmchen F et al (2021) Long-term self-renewing stem cells in the adult mouse hippocampus identified by intravital imaging. Nat Neurosci 24: 225–233

6. Encinas JM, Michurina TV, Peunova N, Park JH, Tordo J, Peterson DA, Fishell G, Koulakov A, Enikolopov G (2011) Division-coupled astrocytic differentiation and age-related depletion of neural stem cells in the adult hippocampus. Cell Stem Cell 8: 566–579

7. Haslinger A, Schwarz TJ, Covic M, Lie DC (2009) Expression of Sox11 in adult neurogenic niches suggests a stage-specific role in adult neurogenesis. Eur J Neurosci 29: 2103–2114

8. Heine VM, Maslam S, Joels M, Lucassen PJ (2004) Prominent decline of newborn cell proliferation, differentiation, and apoptosis in the aging dentate gyrus, in absence of an age-related hypothalamus-pituitary-adrenal axis activation. Neurobiol Aging 25: 361–375

9. Huang da W, Sherman BT, Lempicki RA (2009) Systematic and integrative analysis of large gene lists using DAVID bioinformatics resources. Nat Protoc 4: 44–57

10. Hubner CA, Lorke DE, Hermans-Borgmeyer I (2001) Expression of the Na-K-2Cl-cotransporter NKCC1 during mouse development. Mech Dev 102: 267–269

11. Ibrayeva A, Bay M, Pu E, Jorg DJ, Peng L, Jun H, Zhang N, Aaron D, Lin C, Resler G et al (2021) Early stem cell aging in the mature brain. Cell Stem Cell 28: 955–966 e957

12. Imayoshi I, Ohtsuka T, Metzger D, Chambon P, Kageyama R (2006) Temporal regulation of Cre recombinase activity in neural stem cells. Genesis 44: 233–238

13. Imayoshi I, Sakamoto M, Ohtsuka T, Kageyama R (2009) Continuous neurogenesis in the adult brain. Dev Growth Differ 51: 379–386

14. Kim D, Pertea G, Trapnell C, Pimentel H, Kelley R, Salzberg SL (2013) TopHat2: accurate alignment of transcriptomes in the presence of insertions, deletions and gene fusions. Genome Biol 14: R36

15. Kuhn HG, Dickinson-Anson H, Gage FH (1996) Neurogenesis in the dentate gyrus of the adult rat: age-related decrease of neuronal progenitor proliferation. J Neurosci 16: 2027–2033

16. Kuriakose D, Xiao ZC (2024) Protocol to detect telomerase activity in adult mouse hippocampal neural progenitor cells using the telomeric repeat amplification protocol assay. STAR Protoc 5: 103108

17. Lagace DC, Whitman MC, Noonan MA, Ables JL, DeCarolis NA, Arguello AA, Donovan MH, Fischer SJ, Farnbauch LA, Beech RD et al (2007) Dynamic contribution of nestin-expressing stem cells to adult neurogenesis. J Neurosci 27: 12623–12629

18. Lefebvre V, Dumitriu B, Penzo-Mendez A, Han Y, Pallavi B (2007) Control of cell fate and differentiation by Sry-related high-mobility-group box (Sox) transcription factors. Int J Biochem Cell Biol 39: 2195–2214

19. Liao Y, Smyth GK, Shi W (2014) featureCounts: an efficient general purpose program for assigning sequence reads to genomic features. Bioinformatics 30: 923–930

20. Love MI, Huber W, Anders S (2014) Moderated estimation of fold change and dispersion for RNA-seq data with DESeq2. Genome Biol 15: 550

21. Matsubara S, Matsuda T, Nakashima K (2021) Regulation of Adult Mammalian Neural Stem Cells and Neurogenesis by Cell Extrinsic and Intrinsic Factors. Cells 10

22. Mu L, Berti L, Masserdotti G, Covic M, Michaelidis TM, Doberauer K, Merz K, Rehfeld F, Haslinger A, Wegner M et al (2012) SoxC transcription factors are required for neuronal differentiation in adult hippocampal neurogenesis. J Neurosci 32: 3067–3080

23. Obernier K, Alvarez-Buylla A (2019) Neural stem cells: origin, heterogeneity and regulation in the adult mammalian brain. Development 146

24. Obernier K, Cebrian-Silla A, Thomson M, Parraguez JI, Anderson R, Guinto C, Rodas Rodriguez J, Garcia-Verdugo JM, Alvarez-Buylla A (2018) Adult Neurogenesis Is Sustained by Symmetric Self-Renewal and Differentiation. Cell Stem Cell 22: 221–234 e228

25. Pilz GA, Bottes S, Betizeau M, Jorg DJ, Carta S, April S, Simons BD, Helmchen F, Jessberger S (2018) Live imaging of neurogenesis in the adult mouse hippocampus. Science 359: 658–662

26. Shin J, Berg DA, Zhu Y, Shin JY, Song J, Bonaguidi MA, Enikolopov G, Nauen DW, Christian KM, Ming GL, Song H (2015) Single-Cell RNA-Seq with Waterfall Reveals Molecular Cascades underlying Adult Neurogenesis. Cell Stem Cell 17: 360–372

27. Smith T, Heger A, Sudbery I (2017) UMI-tools: modeling sequencing errors in Unique Molecular Identifiers to improve quantification accuracy. Genome Res 27: 491–499

28. Song J, Zhong C, Bonaguidi MA, Sun GJ, Hsu D, Gu Y, Meletis K, Huang ZJ, Ge S, Enikolopov G et al (2012) Neuronal circuitry mechanism regulating adult quiescent neural stem-cell fate decision. Nature 489: 150–154

29. Urban N, Blomfield IM, Guillemot F (2019) Quiescence of Adult Mammalian Neural Stem Cells: A Highly Regulated Rest. Neuron 104: 834–848

30. Walter J, Keiner S, Witte OW, Redecker C (2011) Age-related effects on hippocampal precursor cell subpopulations and neurogenesis. Neurobiol Aging 32: 1906–1914

31. Wang Y, Lin L, Lai H, Parada LF, Lei L (2013) Transcription factor Sox11 is essential for both embryonic and adult neurogenesis. Dev Dyn 242: 638–653

32. Watanabe M, Fukuda A (2015) Development and regulation of chloride homeostasis in the central nervous system. Front Cell Neurosci 9: 371

33. Woitke F, Blank A, Fleischer AL, Zhang S, Lehmann GM, Broesske J, Haase M, Redecker C, Schmeer CW, Keiner S (2023) Post-Stroke Environmental Enrichment Improves Neurogenesis and Cognitive Function and Reduces the Generation of Aberrant Neurons in the Mouse Hippocampus. Cells 12

34. Wu Y, Bottes S, Fisch R, Zehnder C, Cole JD, Pilz GA, Helmchen F, Simons BD, Jessberger S (2023) Chronic in vivo imaging defines age-dependent alterations of neurogenesis in the mouse hippocampus. Nat Aging 3: 380–390

35. Yamaguchi M, Saito H, Suzuki M, Mori K (2000) Visualization of neurogenesis in the central nervous system using nestin promoter-GFP transgenic mice. Neuroreport 11: 1991–1996

36. Zerbino DR, Achuthan P, Akanni W, Amode MR, Barrell D, Bhai J, Billis K, Cummins C, Gall A, Giron CG et al (2018) Ensembl 2018. Nucleic Acids Res 46: D754–D761

37. Zhang F, Yoon K, Kim NS, Ming GL, Song H (2023) Cell-autonomous and non-cell-autonomous roles of NKCC1 in regulating neural stem cell quiescence in the hippocampal dentate gyrus. Stem Cell Reports 18: 1468–1481

38. Zhang S, Zhou J, Zhang Y, Liu T, Friedel P, Zhuo W, Somasekharan S, Roy K, Zhang L, Liu Y et al (2021) The structural basis of function and regulation of neuronal cotransporters NKCC1 and KCC2. Commun Biol 4: 226

